# Fingerprinting Adiposity and Metabolic Function in the Brains of Overweight and Obese Humans

**DOI:** 10.1101/540997

**Authors:** Michael C. Farruggia, Maria J. Van Kooten, Mary V. Burke, Dustin Scheinost, R. Todd Constable, Dana M. Small

**Affiliations:** Interdepartmental Neuroscience Program, Yale University, 333 Cedar Street, New Haven, CT, U.S; Department of Psychiatry, Yale University School of Medicine, 300 George Street, New Haven, CT 06511, USA; University of Groningen, Faculty of Medical Sciences, Antonius Deusinglaan 1, 9713 AV Groningen, The Netherlands; Department Radiology and Biomedical Imaging, Yale University School of Medicine, New Haven, CT, United States; Department Statistics and Data Science, Yale University, New Haven, CT, United States; Child Study Center, Yale School of Medicine, New Haven, CT, United States; Department of Neurosurgery, Yale University School of Medicine, New Haven, CT, United States; Department of Psychology, Yale University, New Haven, CT, United States; MEG Center, University of Tubingen, Tubingen, Germany

**Author notes:** Correspondence should be addressed to: Dana M Small, PhD (D.M.S.), Michael C Farruggia, MS, (M.C.F.), 1 Church Street, Suite 6a New Haven, CT 06510. Other Contact Information, Maria J. Van Kooten; 1 Church Street, Suite 6a, New Haven, CT, 06510 Mary V. Burke; 1 Church Street, Suite 6a, New Haven, CT, 06510 Dustin Scheinost; 300 Cedar Street, Rm. N131, New Haven, CT, 06519 R. Todd Constable; 300 Cedar Street, Rm. N130, New Haven, CT, 06519. Note: With the exception of its pre-print on bioRxiv, none of the material has been published or is under consideration elsewhere, including the Internet.

**Keywords:** Adiposity, metabolism, waist circumference, insulin, fMRI, functional connectivity, predictive modeling, obesity

## Abstract

The brain plays a central role in the pathophysiology of obesity. Connectome-based Predictive Modeling (CPM) is a newly developed, data-driven approach that exploits whole-brain functional connectivity to predict a behavior or trait that varies across individuals. We used CPM to determine whether brain “fingerprints” evoked during milkshake consumption could be isolated for common measures of adiposity in 67 overweight and obese adults. We found that a CPM could be identified for waist circumference, but not percent body fat or BMI, the most frequently used measures to assess brain correlates of obesity. In an exploratory analysis, we were also able to derive a largely distinct CPM predicting fasting blood insulin. These findings demonstrate that brain network patterns are more tightly coupled to waist circumference than BMI or percent body fat and that adiposity and glucose tolerance are associated with distinct maps, pointing to dissociable central pathophysiological phenotypes for obesity and diabetes.

## Introduction

Obesity is a neurobehavioral disorder resulting from a biologically vulnerable brain that is highly sensitive to its environment *(1)*. Evidence for this derives from genome-wide association studies (GWAS), demonstrating that genetic variants associated with obesity tend to affect brain function. For example, high-risk variants in the FTO (fat mass and obesity associated) gene predispose individuals to weighing on average 3kg more and increasing their obesity risk 1.67x over those with low-risk variants *(2, 3)*. The FTO gene encodes the alpha-ketoglutarate-dependent dioxygenase enzyme, a protein that is highly expressed in the human brain, particularly in the hypothalamus and cerebellum as well as on dopamine receptors where it influences reward learning *(4–6)*. Hypotheses of FTO’s action in the human brain suggest that it may act to increase consumption of food and alter sensing of dietary macronutrient composition *(7)*. Since the discovery of FTO, additional GWAS have found another 97 BMI-associated SNPs which encode for proteins that are significantly enriched in the human brain *(8)*.

While the vast majority of evidence continues to point to the brain as integral in the pathophysiology of obesity, none of the adiposity measures we have to quantify it or characterize its extent are brain measures. Body mass index, or BMI, is the anthropometric measurement commonly used to classify obesity, with individuals having a BMI ≥ 30 characterized as obese *(9)*. However, BMI as a measure of adiposity is often criticized for being an inaccurate reflection of fat mass, in particular in Asians and in athletes *(10, 11)*. Body fat percentage alone has also been criticized as an inaccurate predictor of negative health outcomes, as regional distribution of fat in the body influences risk for negative health consequences. For example, women with predominantly upper body adipose tissue are more likely to have diabetic glucose tolerance results and higher fasting plasma triglycerides than those with predominantly lower body adipose tissue *(12)*. More recent studies have found that distribution of adipose tissue, particularly in the thighs, is inversely correlated with metabolic syndrome in obesity *(13)*.

Ultimately, if obesity is a neurobehavioral disorder resulting from a vulnerable brain interacting with an obesogenic environment, the most informative adiposity measures should be predicted by brain function to some extent. Further, if an adiposity measure can be reliably predicted using features of brain function in a cross-validated framework, it follows that these features may represent a generalizable index of adiposity. Thus, not only is it important that we are able to determine which adiposity measure is best predicted by whole-brain functional connectivity, but it also imperative to determine which anatomical regions (i.e. nodes) and which connections among them (i.e. edges) are influencing our predictive power.

With this in mind, we set out to determine which commonly-used adiposity measures, including BMI, body fat %, waist size, body weight, and waist/hip ratio, could be predicted by whole-brain function connectivity. To accomplish this, we use connectome-based predictive modeling (CPM), a data-driven approach that builds predictive models of brain-behavior relationships using connectivity data in a cross-validated framework *(14, 15)*. More specifically, CPM exploits individual variation in whole-brain functional connectivity to predict a behavior or trait that varies across individuals in a leave-one-out procedure. CPM has been used to identify brain “fingerprints” *(14)* as well as to predict personality traits *(16)*, creative ability *(17)*, and sustained attention *(18)*. Here we used this technique to ask whether a predictive CPM could be identified for our different adiposity measures. As an exploratory aim, we also tested whether a CPM could be found for measures of glucose tolerance.

## Results

### CPM predicts individual differences in waist circumference

Of all measures, including BMI, body fat %, waist size, body weight, and waist/hip ratio, we found that CPM significantly predicts waist circumference (combined network, r = 0.354, p = 0.005) after applying Benjamini-Hochberg FDR correction for multiple tests (Fig. 1). Overall, the combined model accounts for approximately 12.5% of the variance in the waist circumference of a novel subject.

**Fig. 1.**
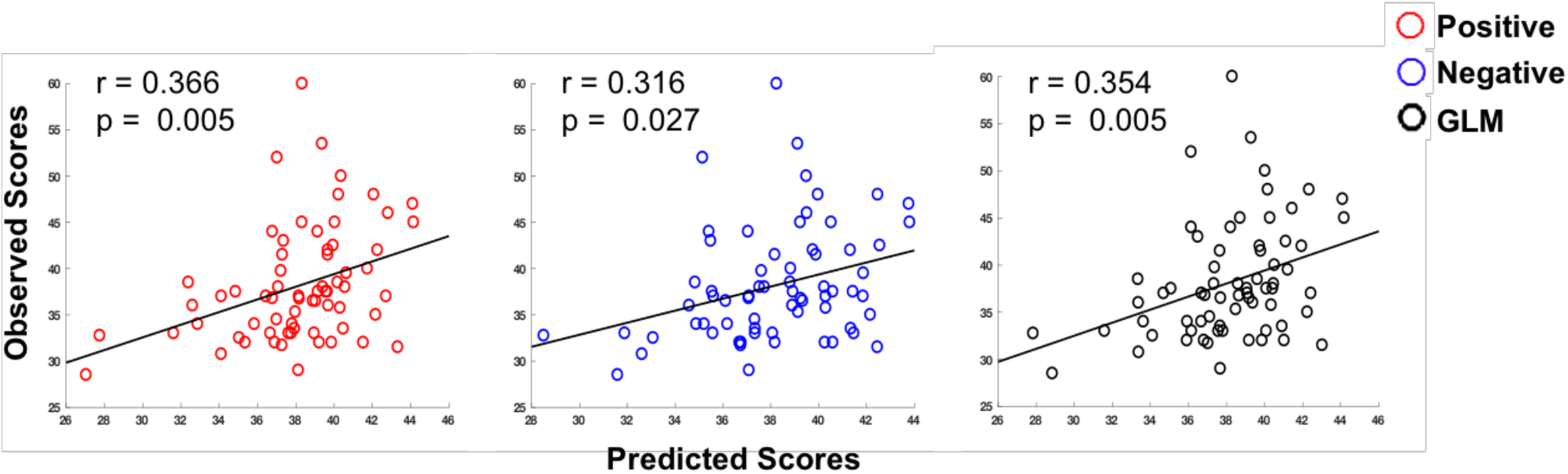
Relationship Between Observed and Predicted Waist Circumference. Scatterplots relating observed versus predicted waist circumference in positive, negative, and combined networks. Positive networks consist of edges that are positively correlated with behavior, while negative networks are constructed using edges that are negatively correlated with behavior. Combined networks are constructed using a general linear model (GLM). Overall, approximately 13% of the variance in a novel individual’s waist circumference can be accounted for using CPM.

### Waist circumference CPM contains DMN, motor, visual, cerebellar, and subcortical networks

The combined model consists of edges whose strengths are positively (positive network) or negatively (negative network) correlated with our variable of interest, waist circumference. These edges can be visualized as connections among nodes assigned to one of ten bilateral brain regions (Fig. 2A), or as the density of edges among ten canonical networks normalized by network size (Fig. 2B). Brain regions include Prefrontal, Motor, Insula, Parietal, Temporal, Occipital, Limbic, Cerebellum, Subcortical, and Brainstem. We operationally define our high degree nodes as the top N nodes that occur before multiple degree repetitions. High degree nodes (degree ≥ 9), or nodes with the highest number of connections for the waist circumference CPM were found in prefrontal, parietal, limbic, cerebellar, subcortical, and temporal regions (Fig. 2A, *see also 5A*). AAL labels for these regions include left inferior frontal gyrus, left postcentral gyrus, left amygdala, left culmen of cerebellum, right putamen and right fusiform gyrus, respectively (Table 1).

**Fig. 2.**
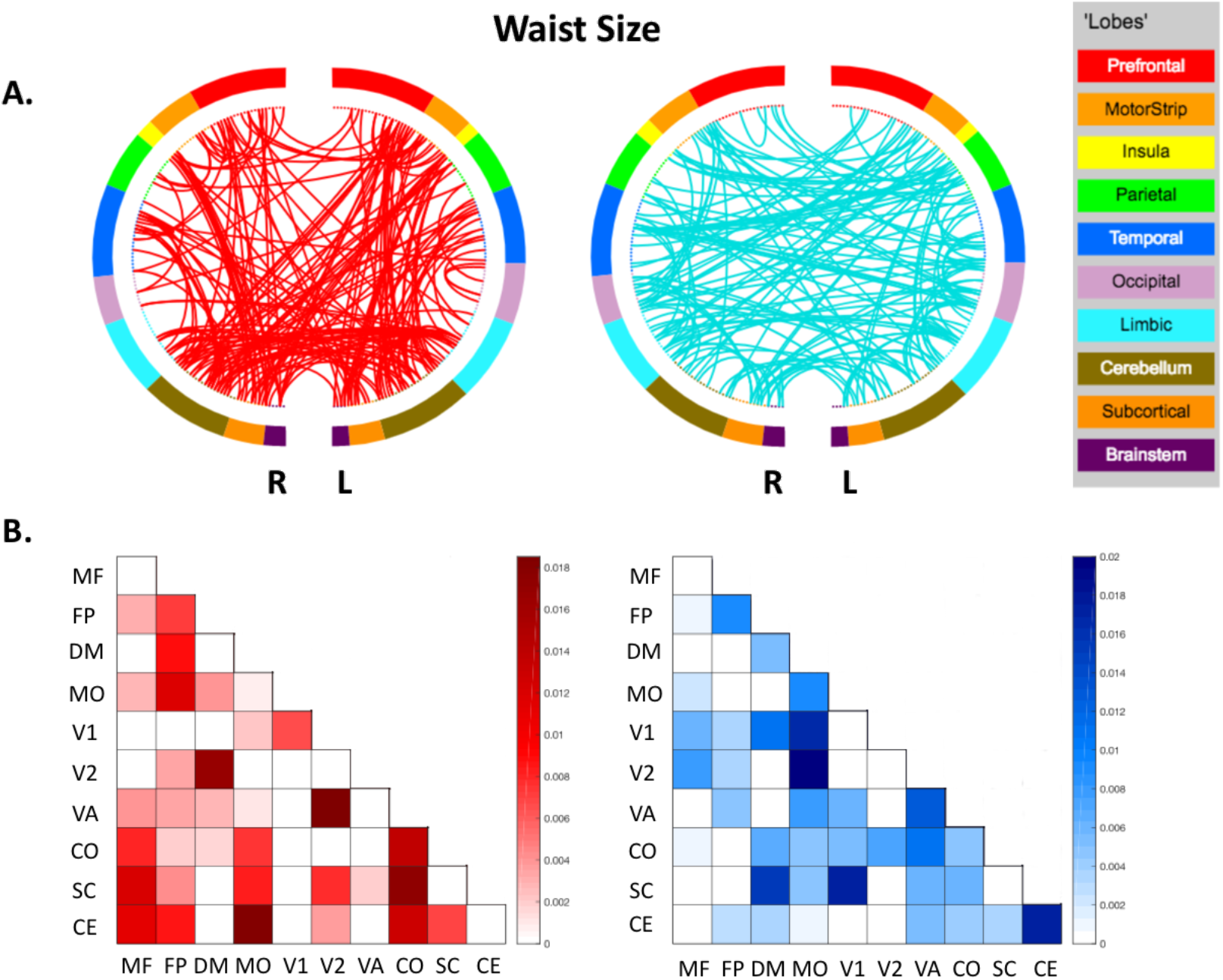
Visualization of Positive and Negative Predictive Networks. Representations of positive (red) and negative (blue) predictive networks in **(A)**. Each semicircle is a different hemisphere of the brain. Connections represent edges between nodes, with nodes organized among ten macroscale regions. Matrices **(B)** show the proportion of positive versus negative edges shared between network pairs among ten canonical networks normalized for network size (MF: medial frontal; FP: frontoparietal; DM: default mode; MO: Motor; V1: visual 1; V2: visual 2; VA: visual association; CO: cingulo-opercular; SC: subcortical; CE: cerebellar).

**Table 1.**
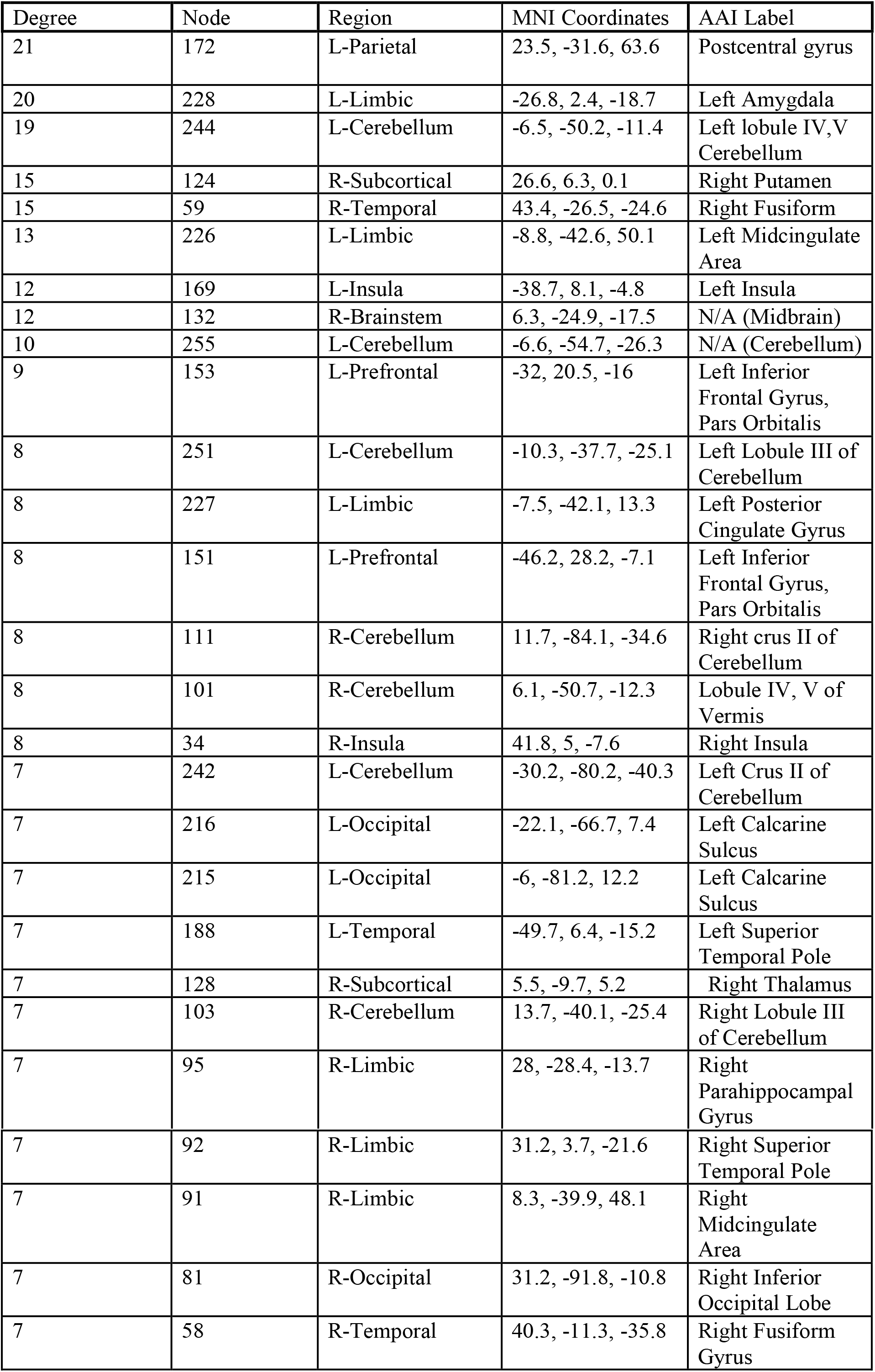
Top 10% High-Degree Nodes in the Waist Circumference Network.

| Degree | Node | Region | MNI Coordinates | AAI Label |
| --- | --- | --- | --- | --- |
| 21 | 172 | L-Parietal | 23.5, -31.6, 63.6 | Postcentral gyrus |
| 20 | 228 | L-Limbic | -26.8, 2.4, -18.7 | Left Amygdala |
| 19 | 244 | L-Cerebellum | -6.5, -50.2, -11.4 | Left lobule IV,V<br>Cerebellum |
| 15 | 124 | R-Subcortical | 26.6, 6.3, 0.1 | Right Putamen |
| 15 | 59 | R-Temporal | 43.4, -26.5, -24.6 | Right Fusiform |
| 13 | 226 | L-Limbic | -8.8, -42.6, 50.1 | Left Midcingulate<br>Area |
| 12 | 169 | L-Insula | -38.7, 8.1, -4.8 | Left Insula |
| 12 | 132 | R-Brainstem | 6.3, -24.9, -17.5 | N/A (Midbrain) |
| 10 | 255 | L-Cerebellum | -6.6, -54.7, -26.3 | N/A (Cerebellum) |
| 9 | 153 | L-Prefrontal | -32, 20.5, -16 | Left Inferior<br>Frontal Gyrus,<br>Pars Orbitalis |
| 8 | 251 | L-Cerebellum | -10.3, -37.7, -25.1 | Left Lobule III of<br>Cerebellum |
| 8 | 227 | L-Limbic | -7.5, -42.1, 13.3 | Left Posterior<br>Cingulate Gyrus |
| 8 | 151 | L-Prefrontal | -46.2, 28.2, -7.1 | Left Inferior<br>Frontal Gyrus,<br>Pars Orbitalis |
| 8 | 111 | R-Cerebellum | 11.7, -84.1, -34.6 | Right crus II of<br>Cerebellum |
| 8 | 101 | R-Cerebellum | 6.1, -50.7, -12.3 | Lobule IV, V of<br>Vermis |
| 8 | 34 | R-Insula | 41.8, 5, -7.6 | Right Insula |
| 7 | 242 | L-Cerebellum | -30.2, -80.2, -40.3 | Left Crus II of<br>Cerebellum |
| 7 | 216 | L-Occipital | -22.1, -66.7, 7.4 | Left Calcarine<br>Sulcus |
| 7 | 215 | L-Occipital | -6, -81.2, 12.2 | Left Calcarine<br>Sulcus |
| 7 | 188 | L-Temporal | -49.7, 6.4, -15.2 | Left Superior<br>Temporal Pole |
| 7 | 128 | R-Subcortical | 5.5, -9.7, 5.2 | Right Thalamus |
| 7 | 103 | R-Cerebellum | 13.7, -40.1, -25.4 | Right Lobule III<br>of Cerebellum |
| 7 | 95 | R-Limbic | 28, -28.4, -13.7 | Right<br>Parahippocampal<br>Gyrus |
| 7 | 92 | R-Limbic | 31.2, 3.7, -21.6 | Right Superior Temporal Pole |
| 7 | 91 | R-Limbic | 8.3, -39.9, 48.1 | Right Midcingulate Area |
| 7 | 81 | R-Occipital | 31.2, -91.8, -10.8 | Right Inferior Occipital Lobe |
| 7 | 58 | R-Temporal | 40.3, -11.3, -35.8 | Right Fusiform Gyrus |

Canonical networks emphasize different functional properties of the human brain and consist of medial frontal (MF), frontoparietal (FP), default mode (DM), motor (MO), visual 1 (V1), visual 2 (V2), visual association (VA), cingulo-opercular (CO), subcortical (SC), and cerebellum (CE) networks *(19, 20)*. The highest proportion of edges, that is, the edges used most in prediction for the positive network exist between DM-V2, MO-CE, V2-VA, and CO-SC. In the negative network, these are between DM-SC, MO-V2, MO-V1, V1-SC, VA-VA, and CE-CE (Fig. 2B).

### Exploratory analysis: CPM predicts individual differences in fasting insulin

Due to the large scope of the dataset, we were able to conduct a post-hoc exploratory analysis of metabolic markers to examine convergence or divergence of predictive networks as they relate to the waist circumference CPM. For this analysis, we analyzed additional bloods data until we had a complete metabolic panel from all 59 of 67 subjects. Fasting glucose, fasting insulin, and HOMA-IR were input into CPM as our measures of interest. CPM was able to predict fasting insulin (p < 0.05, uncorrected; Fig. 3A) but not fasting glucose or HOMA-IR, accounting for approximately 7% of the variance in a novel individual’s fasting insulin levels. Positive and negative predictive networks for fasting insulin are notably sparser than those for waist circumference (Fig. 3B). High degree nodes (degree ≥ 6) for these networks were found in cerebellar (left crus and right lobule VIIB), temporal (left superior temporal pole and right fusiform), and limbic (right posterior cingulate gyrus) regions (Fig. 3B, *see also 5B, Table 2*). In the positive network, a high density of edges exists between MF-CE, DM-V2, MO-CE, V2-CE, and V2-SC (Fig. 3C). In the negative network, these exist between MF-MF, FP-VA, FP-V1, DM-CE, DM-SC, V1-SC, V1-V2, VA-CO, CO-CE, and SC-SC (Fig. 3C).

**Fig. 3.**
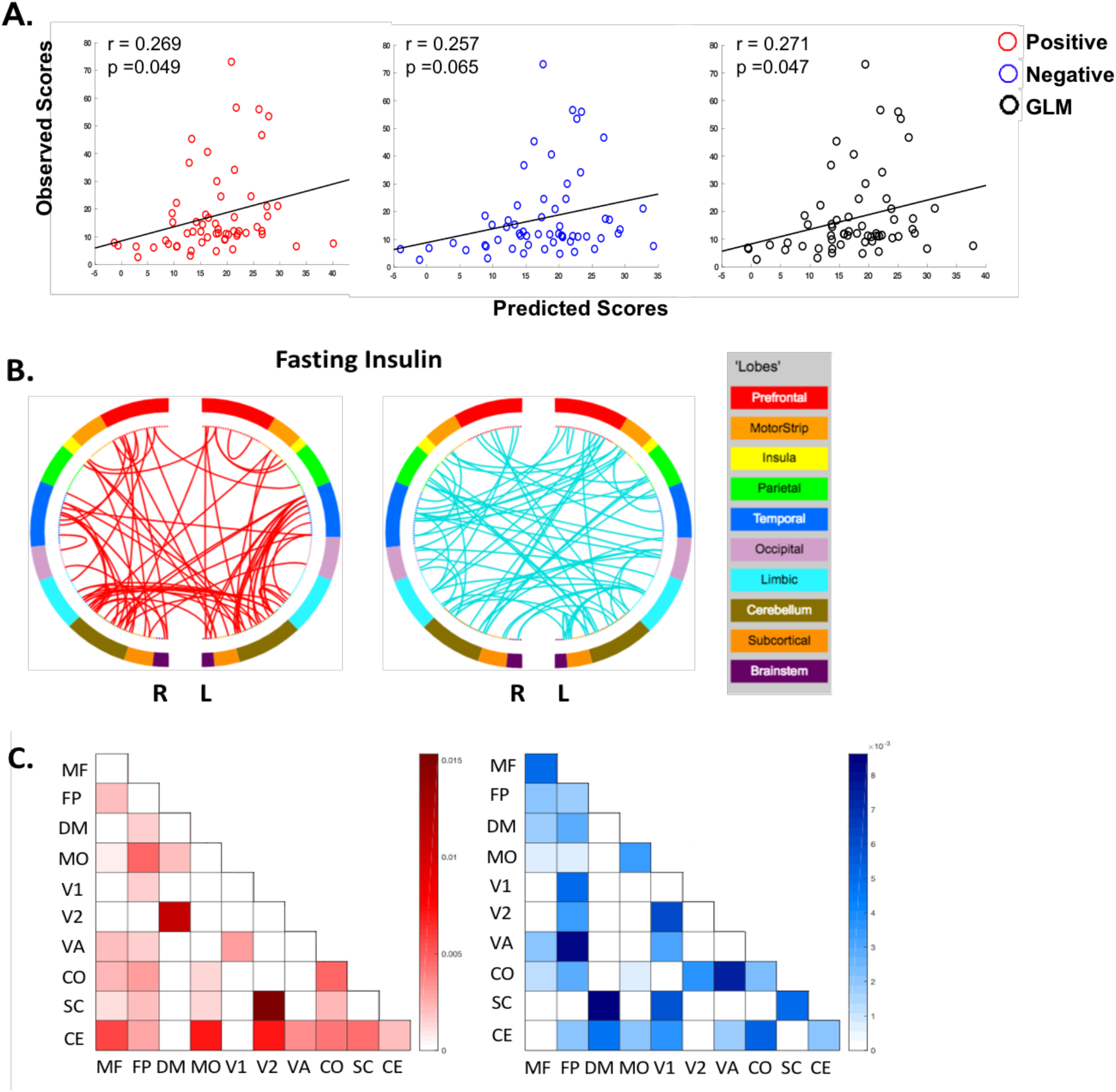
Exploratory Analysis of Fasting Insulin. CPM predicts fasting insulin levels **(A)**. Positive and negative predictive networks are shown as hemisphere plots in **(B)**. Notably, the number of positive and negative edges are highest among cerebellar and frontoparietal regions and the rest of the brain **(C)**.

**Table 2.** Top 10% High-Degree Nodes in the Fasting Insulin Network.

| Degree | Node | Region | MNI Coordinates | AAL Label |
| --- | --- | --- | --- | --- |
| 11 | 238 | L-Cerebellum | -37, -52.9, -31.1 | Left Crus I of Cerebellum |
| 8 | 186 | L-Temporal | -34.6, 18.6, -32.2 | Left Superior Temporal Pole |
| 7 | 227 | L-Limbic | -7.5, -42.1, 13.3 | Left Posterior Cingulate Gyrus |
| 7 | 59 | R-Temporal | 43.4, -26.5, -24.6 | Right Fusiform |
| 6 | 116 | R-Cerebellum | 41.9, -64, -49.2 | Right Lobule VIIB of Cerebellum |
| 5 | 103 | R=Cerebellum | 13.7, -40.1, -25.4 | Right Lobule III of Cerebellum |
| 5 | 102 | R-Cerebellum | 39.1, -74.9, -29.7 | Right Crus I of Cerebellum |
| 5 | 92 | R-Limbic | 31.2, 3.7, -21.6 | Right Superior Temporal Pole |
| 4 | 265 | L-Brainstem | -5, -21.5, -15.8 | N/A (Midbrain) |
| 4 | 263 | L-Subcortical | -4.9, -10.3, 5.8 | Left Thalamus |
| 4 | 208 | L-Occipital | -16.8, -84.9, 33 | Left Cuneus |
| 4 | 188 | L-Temporal | -49.7, 6.4, -15.2 | Left Superior Temporal Pole |
| 4 | 154 | L-Prefrontal | -43, 42, 11 | Left Inferior Frontal Gyrus Pars Triangularis |
| 4 | 139 | L-Prefrontal | -18.2, 57, -14.3 | Left Superior Frontal Gyrus, Orbital part |
| 4 | 99 | R-Limbic | 19.3, -7.7, -14.8 | Right Hippocampus |
| 4 | 36 | R-Insula | 37.5, 21.1, -10.1 | Right Inferior Frontal Gyrus, Orbital part |
| 3 | 255 | L-Cerebellum | -6.6, -54.7, -26.3 | N/A (Cerebellum) |
| 3 | 254 | L-Cerebellum | -21.2, -53.4, -23.6 | Left Lobule VI of Cerebellum |
| 3 | 253 | L-Cerebellum | -26.3, -69.5, -30.6 | Left Crus I of Cerebellum |
| 3 | 252 | L-Cerebellum | -46.4, -46.8, -42.9 | Left Crus II of Cerebellum |
| 3 | 251 | L-Cerebellum | -10.3, -37.7, -25.1 | Left lobule III of Cerebellum |
| 3 | 246 | L-Cerebellum | -42.6, -63.7, -46.3 | Left Crus II of Cerebellum |
| 3 | 242 | L-Cerebellum | -30.2, -80.2, -40.3 | Left Crus II of Cerebellum |
| 3 | 202 | L-Temporal | -30, -5.8, -40.9 | Left Fusiform Gyrus |
| 3 | 163 | L-MotorStrip | -57, -3.4, 6.8 | Left Superior Temporal Gyrus |
| 3 | 135 | L-Prefrontal | -18.2, 19, -21 | Left Inferior Frontal Gyrus, Pars Orbitalis |
| 3 | 132 | R-Brainstem | 6.3, -24.9, -17.5 | N/A (Midbrain) |

### Waist circumference and fasting insulin CPMs are separable

To examine convergence and divergence of predictive networks, we subtracted the number of negative edges from the number of positive edges between each canonical network and plotted the resultant degree of polarity (Fig. 4A-B). For example, networks can have mostly positive or negative edges connecting them (i.e. polarity), or an even distribution of positive and negative edges (i.e. apolarity). Indeed, this technique demonstrates the degree to which positive or negative edges are used in prediction and can help ascertain common or uncommon neurobiological correlates between fasting insulin and waist circumference. The predictive network for waist size shows a large degree of polarity (Fig. 4A), while the relative sparsity of the predictive network for insulin is visible and there is notable apolarity particularly among visual networks (Fig. 4B). Predictive networks for insulin and waist size are on average dissimilar. Of the 55 network pairs, 26 network pairs (47%) exhibited similar degrees of polarity between the waist circumference and insulin predictive networks. Most notably, patterns of connectivity within cerebellar and subcortical networks showed the most similar patterns of polarity between waist circumference and fasting insulin predictive networks (Fig. 4). In contrast their polarity differs most in medial-frontal, frontoparietal, default mode, motor, visual, and cingulo-opercular networks. To aid in comparison, we plotted the spatial extent of the top 10% highest-degree predictive nodes for each network (Fig. 5A-B, Tables 1 & 2). Only nine of the top 10% of nodes (27 total) in each CPM are shared among each other, those being centered in left posterior cingulate gyrus (node 227), right fusiform (59), right and left superior temporal pole (92 and 188), left cerebellum (255), left and right lobule III of cerebellum (251 and 103), left crus II of cerebellum (242), and midbrain (132). The 18 nodes (66%) that differ are depicted in Fig. 6A-B.

**Fig. 4.**
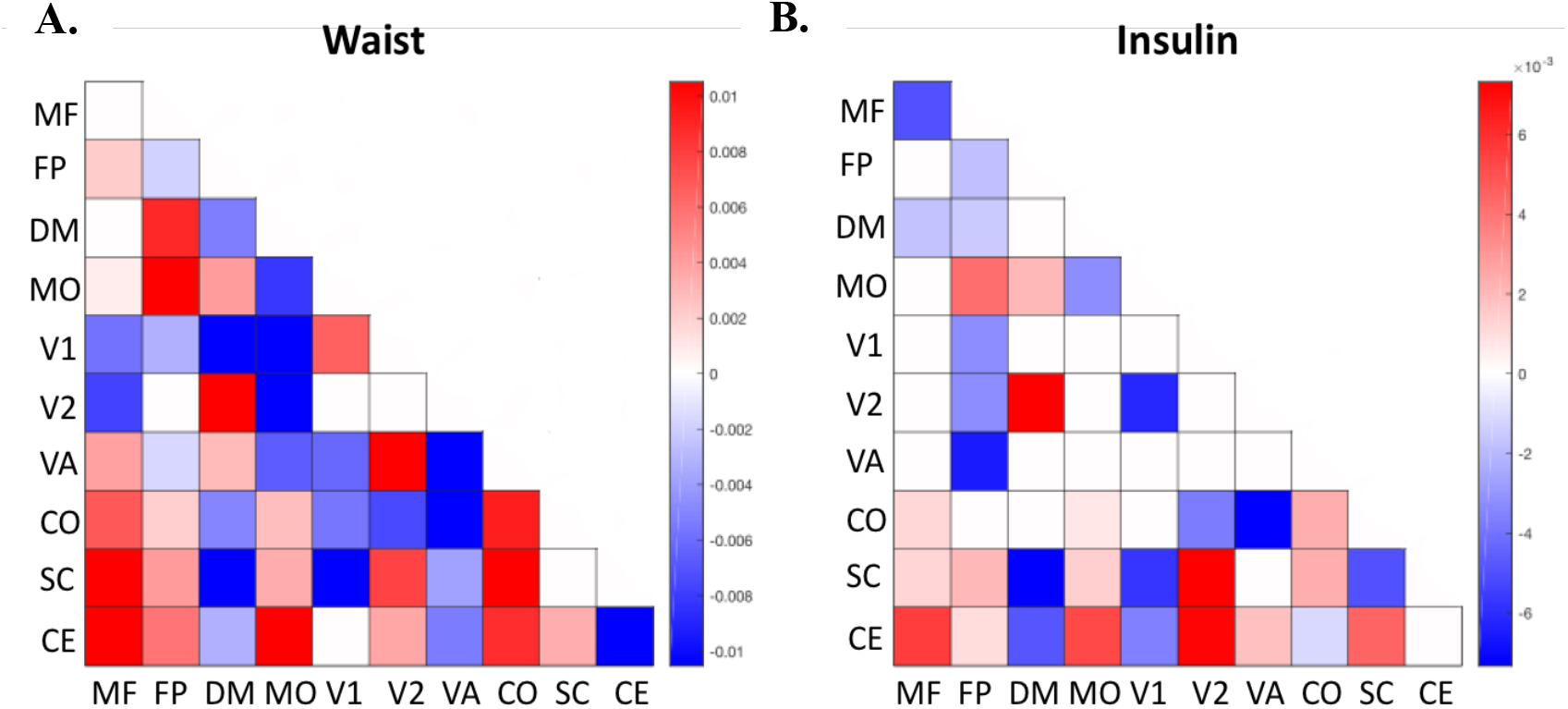
Difference Matrices of Waist Circumference and Insulin. Matrices express the polarity in the number of positive versus negative edges between canonical networks adjusted for network size. For example, **(A)** shows more edges in the positive network exist between cerebellar and medial frontal nodes, while more edges in the negative network exist for cerebellar-cerebellar connections. Taken together, **(A)** and **(B)** demonstrate the greatest degree of similarity in connectivity patterns of cerebellar and subcortical networks in waist circumference and fasting insulin CPMs.

**Fig. 5.**
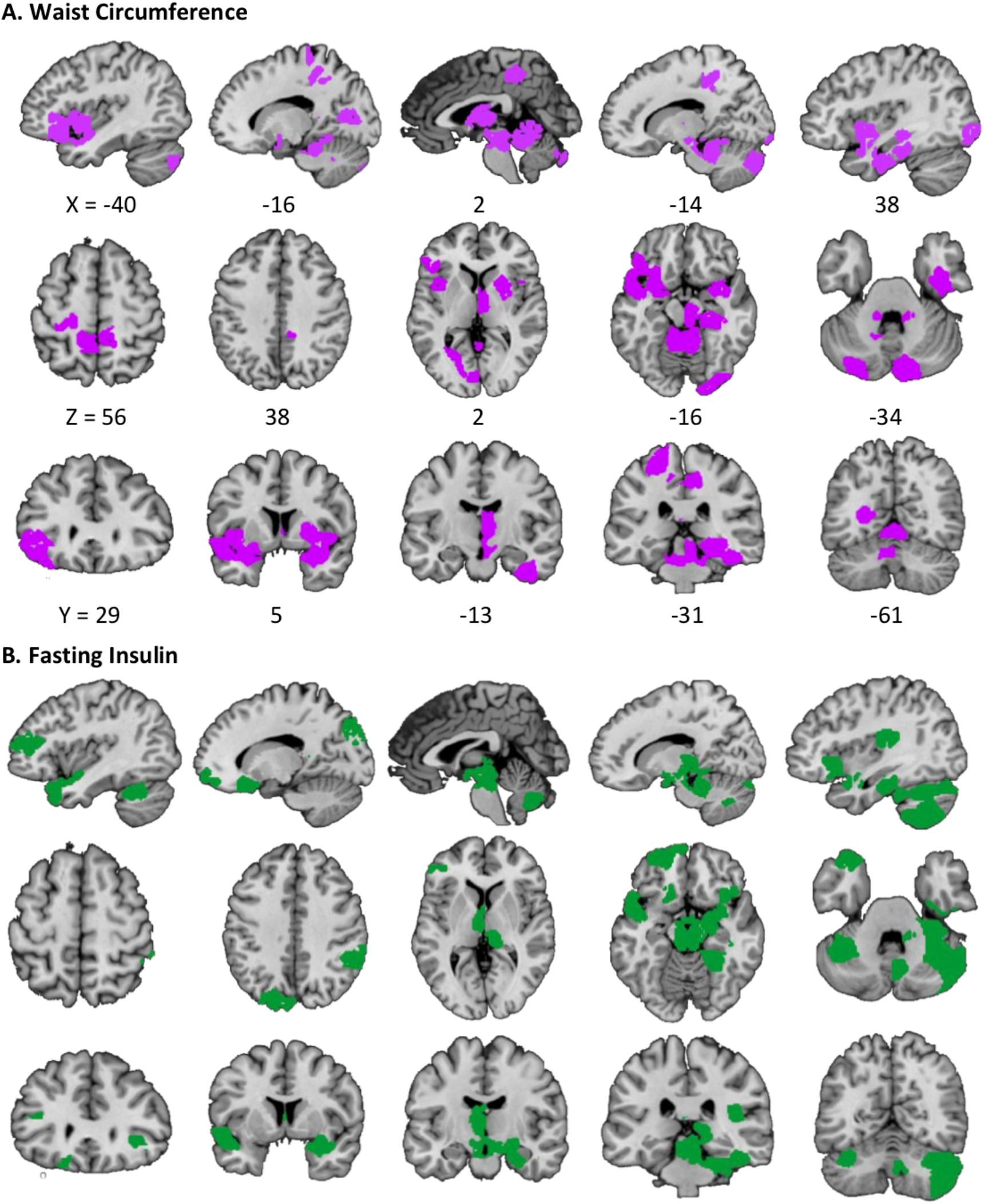
Top 10% High-Degree Nodes for Waist Circumference and Fasting Insulin CPMs. Top 10%) high-degree nodes (27 nodes total) for waist circumference **(A)** encompass networks spanning the whole brain and include nodes varying in degree from 21 to 7. **(B)** The CPM for fasting insulin is similarly encompassing, with nodes varying in degree from 11 to 3.

**Fig. 6.**
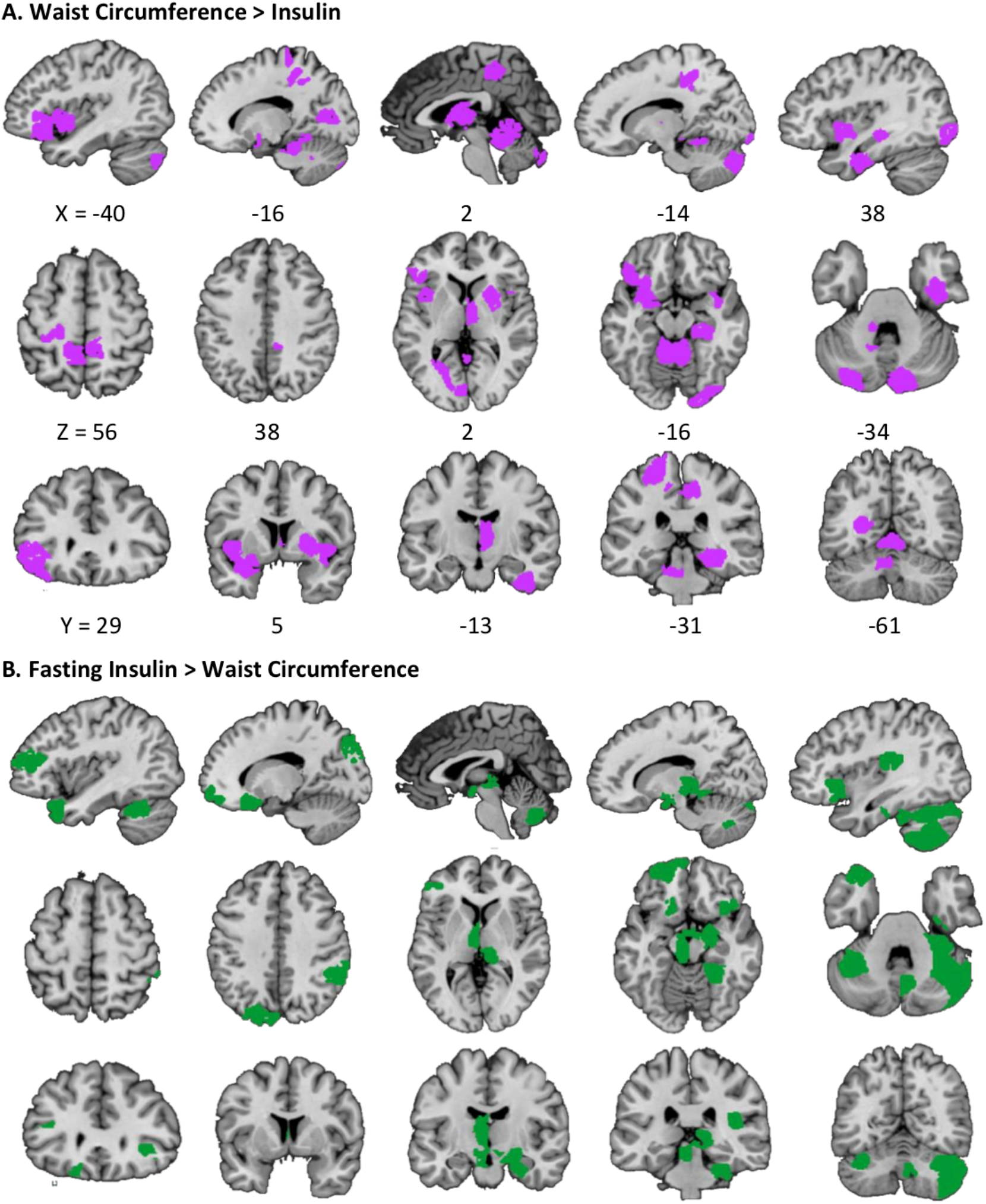
Difference of Top 10% Nodes between Waist Circumference and Fasting Insulin CPMs. **(A)** High-degree nodes (top 10%) participating in the waist circumference CPM but not the fasting insulin CPM **(B)** Nodes in the top 10% participating in the fasting insulin CPM but not the waist circumference CPM.

### The role of subject motion

We used frame-to-frame displacement (FTF) as a gross measure of head motion *(21)*. FTF was calculated for each run in each subject and averaged across runs. FTF significantly correlated with waist size (R^2^ = 0.199, p < 0.001), BMI (R^2^ = 0.115, p < 0.01), body weight (R^2^ = 0.132, p < 0.01), waist/hip ratio (R^2^ = 0.103, p < 0.01), and BF% (R^2^ = 0.107, p < 0.01). FTF was notably not correlated with fasting glucose (r = 0.05, p = 0.68), fasting insulin (r = 0.15, p = 0.25), or HOMA-IR (r = 0.15, p = 0.27). While a correlation between any of our adiposity measures and FTF suggests that motion should be accounted for using partial Pearson correlation, recent evidence suggests that in-scanner head motion is heritable and phenotypically correlated with BMI (22). Indeed, if head motion and adiposity contain shared genetic influences, then controlling for motion in CPM would unnecessarily reduce its predictive power in this instance. Instead we corrected for motion during preprocessing (see Methods) and report results following partial Pearson correlation for mean FTF subject (see Supplementary Table 2). Of note, CPMs for both waist circumference and insulin were both significant, but did not survive FDR correction when motion is included as a covariate.

## Discussion

Our experiment produced several novel findings. First, we determined, using a data-driven approach, that waist circumference could be predicted by whole-brain functional connectivity. In particular, with CPM we can account for approximately 12.5% of the variance in this measure in a novel individual. Notably, we could not predict BMI, the most commonly used measure to define obesity, or other adiposity measures, such as body fat %, body weight, and waist/hip ratio. Second, we found that a metabolic marker, fasting insulin, could be predicted from whole-brain functional connectivity and that this predictive network was distinct from that of waist circumference.

There is long-standing controversy regarding the optimal method to quantify adiposity. We assert that, not only is it important to use a measure that captures an individual’s risk for morbidity and mortality, but it is equally important to capture obesity’s underpinnings as a neurobehavioral disorder. Waist circumference does both.

This proposal is consistent with a number of recent findings. Waist circumference has better discriminatory capability for type 2 diabetes than either BMI or body adiposity index [BAI, calculated as (hip circumference)/((height)^1.5^)–18] in a northern Iranian population *(23, 24)*. A systematic review found that waist circumference performed 3% better than BMI in discriminating adults with hypertension, T2D, metabolic syndrome, dyslipidaemia, and general cardiovascular outcomes *(25)*. A meta-analysis of some 259,200 individuals found that high waist circumference is a better predictor than high BMI for the development of diabetes across all ethnicities tested and for the development of hypertension among Hispanics/Latinos in particular *(26)*. Additional evidence suggests that individuals who are classified as obese by waist circumference have a greater risk for all-cause and cardiovascular disease-related death than individuals who are classified as obese by BMI *(27)*. In a separate cohort of 2266 hypertensive patients, waist circumference was able to predict the development of coronary artery disease, while other measures of adiposity such as BMI and waist/hip ratio had no independent prognostic value *(28)*.

Although most neuroimaging studies use BMI to assess neural correlates of obesity, when waist circumference is pitted against BMI, stronger and more extensive correlations are often found. For example, waist circumference, is a better predictor of local gray matter volume *(29, 30)* and brain response to food cues *(31–33)*. The current study confirms and extends these findings by using functional connectivity and a whole-brain data-driven approach to show that brain measures not only correlate most strongly with waist circumference, but also are able to predict waist circumference. This indicates that this neural measure may act as a biomarker for the measure of adiposity that best predicts obesity-related health outcomes *(34)*.

More specifically, high-degree nodes for the waist circumference CPM were located in left inferior frontal gyrus, left postcentral gyrus, left amygdala, the left culmen of the cerebellum, right putamen, and right fusiform gyrus (Fig. 5A). Enhanced responses to food-related stimuli can be found in many of these regions and responses in some regions have been associated with weight gain *(35, 36)*. However, in these instances the relationship is correlative. For example, amygdala response to food related stimuli when sated, but not hungry, correlates with 12-month weight gain *(37)*. This suggests that amygdala responsivity to food cues in the absence of hunger is predictive of overeating and weight gain, but it is not a direct test of the hypothesis because the amygdala response could be epiphenomenal. Likewise, response in the right putamen, during milkshake consumption is predictive of weight change with direction depending upon genotype *(38)*. The current results go beyond these findings. CPM presents as a strength its cross-validated framework based on data-driven model optimization. Here we were able to identify brain features that predict and therefore may constitute a neural index of waist circumference.

This result raises the question as to how the CPM we have identified contributes to the development of increased waist circumference and conversely, how and why waist circumference specifically influences the identified brain “fingerprint”. Notably, although waist circumference is the best adiposity predictor of glucose control, inspection of the anatomy of the CPMs identified for waist circumference and fasting insulin, a marker of insulin resistance, reveals that they are largely distinct (Fig. 5, 6) *(39 –41)*. In particular, the CPMs for waist circumference and fasting insulin showed minimal overlap in the highest degree nodes. This suggests there are distinct neuropathophysiological features. The distinction between the CPMs is best captured by their relative polarity, or the number of positive versus negative edges between networks. The cerebellar and subcortical networks show similar patterns of connectivity while patterns in medial-frontal, frontoparietal, default mode, motor, visual, and cingulo-opercular networks largely differ. Overlap between cerebellar and subcortical networks for adiposity and metabolic dysfunction suggests that these regions may play an integral role in the pathophysiology of obesity. These observations support future efforts to uncover the link between the distinct networks and their contributions to the causes and consequences of obesity. Our findings also have clinical implications. One important application of neuroimaging in clinical practice, is to enable the prediction of clinical outcomes using brain structure, activity, or connectivity *(42)*. Previous methods in predicting brain-behavior relationships tended to be correlational, and thus would overfit to data and fail to generalize to novel individuals *(15)*. As indicated above, CPM presents as a strength its cross-validated framework, which suggests that the predictive features for waist circumference and fasting insulin are generalizable within novel subjects in this dataset. In other words, it is a window into the brain features that contribute to predicting adiposity and metabolic outcomes and as such may constitute biomarkers to index risk or the efficacy of therapeutic interventions.

Further, to apply such a finding to clinical practice, network-based prediction models of obesity must be shown to generalize across samples, populations, and experimental settings *(42)*. A promising avenue to then implement CPM as a tool for clinical use is to use real-time fMRI biofeedback to change connectivity patterns or activation profiles within predictive networks across subjects. Similar approaches have been successfully used in previous studies, namely those attempting to alleviate contamination anxiety, enhance emotion regulation capabilities, and mitigate auditory verbal hallucinations in schizophrenia, among others *(43–45)*. Another approach, similar to that of Rosenberg et al., 2016 *(46)*, is to treat overweight and obesity using a particular intervention and observe how the predictive networks change. We predict that more efficacious treatments will reduce network strength within networks that predict high waist circumference and fasting insulin, and increase the strength of antithetical networks. Recent work suggests that networks obtained using CPM can predict abstinence during treatment for cocaine dependence *(47)*. If the obtained CPMs are relevant to the pathophysiology of obesity, it is thus possible that they may predict weight loss during an intervention, which would be useful for obese individuals seeking treatment. Finally, not only is waist circumference better than BMI and other adiposity measures at predicting a variety of adverse health-related outcomes, but it is also better predicted from whole-brain functional connectivity using CPM. This suggests that including waist circumference as a measure of interest in a clinical setting is imperative to accurately quantify the extent and progression of obesity.

### Limitations and future directions

This study has a number of limitations to address. First, the sample size was relatively limited for a study implementing CPM. It is quite possible that with more data we would be able to build a more informative predictive model. Second, we only include overweight and obese individuals in our dataset due to the nature and design of the longitudinal study for which this data was originally collected. It is thus possible that these CPMs for waist circumference and fasting insulin do not generalize to lean individuals. However, since biomarkers are needed to evaluate therapeutic interventions it can be argued that an overweight/obese sample is most appropriate. Third, because the study design was not specifically optimized for use in this analysis, it remains to be seen whether inclusion of tasteless blocks reduced our predictive power. Tasteless may be more similar to a classic rest condition than milkshake. Across all studies implementing CPM, resting state is least able to maximize detection of individual differences *(19)*. Fourth, we did not use DEXA or whole-body MRI, which are considered the optimal measurement devices to determine adiposity *(48)*. Although, waist circumference and fasting insulin are more practical measures, future work should validate our observations using the gold standard measures. Finally, longitudinal studies are necessary to determine whether the obtained predictive networks arise as a cause or consequence of increased fasting insulin and waist circumference.

## Conclusions

Our analyses reveal that by characterizing whole brain connectivity during consumption of a palatable energy-dense milkshake it is possible to predict an individual’s waist circumference but not BMI or percent body fat. This finding suggests that waist circumference is the phenotypic marker of obesity that is most closely related to brain function during food consumption. However, whether this brain biomarker is a cause or a consequence of waist size is unknown. We were also able to predict fasting insulin but not glucose or HOMA-IR. Notably, although fasting insulin reflects insulin resistance *(49)* and waist circumference has been shown to correlate with insulin resistance, the CPMs predictive of waist circumference and fasting insulin were largely distinct.

## Materials and Methods

### Participants

We recruited 100 overweight and obese individuals using media, flyer advertisements, and the Yale Center for Clinical Investigation Help Us Discover database. Subjects were recruited to participate in an ongoing clinical trial (NCT 01976156), but all related fMRI scans occurred at baseline prior to any experimental manipulations. We screened participants via phone or online to ensure they met the following criteria: right-handed, English-speaking, 18-45 years of age, body mass index (BMI) of 25 or higher, nonsmokers, no psychiatric or neurological impairments (including drug addiction, eating disorders), no obesity-related diseases such as type-2 diabetes, no food allergies, no chemosensory impairments, and no MRI contraindications (pregnancy, claustrophobia, metal implants, or prior head injury with loss of consciousness). In total, 67 participants (mean age = 30.7yrs, mean BMI = 33.3 ± 6.5) met these criteria and underwent fMRI procedures. All participants provided written informed consent according to our protocol that was approved by the Yale University School of Medicine Human Investigation Committee.

### Measures

This study was part of a larger clinical trial (NCT 01976156). Below we outline the procedures related to the baseline fMRI assessment, on which the analyses in this study are based.

### Physical Characteristics and Diet

Participants arrived at the lab on the week of fMRI scanning for anthropometric measurements, which included body weight, height, waist size, hip size, and adiposity. Adiposity, as described by body fat percentage (BF%), was measured using air displacement plethysmography (BodPod). Waist/Hip ratio and BMI (kg/cm^2^) were subsequently calculated. Additional participant characteristics are shown in Supplementary Table 1.

### Bloods

Following an overnight fast of 10-12 hours, blood samples were collected from 59 out of 67 subjects. We were unable to collect bloods on eight subjects due to venipuncture difficulty. Samples were collected, centrifuged immediately, and then frozen at either 80 °C (insulin) or 20 °C (glucose). Insulin protein concentrations were subsequently measured using a radioimmunoassay kit (APLCO insulin ELISA Catalog #80-INSHU-E01.1, Salem, NH) and plasma glucose concentrations were determined using a glucose oxidase analyzer (YSI Incorporated, Yellow Springs, OH). Insulin sensitivity was calculated using the homeostasis model of insulin resistance, or HOMA-IR, which is based on the following formula: [insulin (mU/L) x glucose (mmol/L)]/22.5 *(50)*.

### Image protocols

All fMRI scans were scheduled during the week of blood draws, sometime between 8:00 AM and 12:00 PM. Participants were instructed to arrive neither full nor hungry, and adherence to these instructions were confirmed using internal state ratings. Structural and functional imaging data were then collected on a Siemens 3.0T TIM Trio scanner at the Yale Magnetic Resonance Research Center (MRRC), using the following scan parameters: high resolution T1-weighted structural scan repetition time (TR) = 2530ms, echo time (TE) = 3.31ms, slice thickness = 1mm, flip angle = 7°, field of view (FOV) = 256 x 256, 256 x 256 matrix, 176 slices; functional milkshake run TR = 2000ms, TE = 20 ms, slice thickness = 3mm, flip angle = 80°, FOV = 220 x 220, matrix = 64 x 64, 40 contiguous slices.

fMRI scans consisted of two 9-minute runs that were divided into blocks of 4-8 cued presentations of milkshake or tasteless solutions. These solutions were delivered in quantities of 0.5 mL over a 2s period followed by 7s to swallow. Different milkshake flavors were used to reduce the possibility of stimulus habituation. After each block, participants rinsed their mouths with 0.5mL of deionized water over a period of 2s and were given 3s to swallow. Blocks varied in length from 36s to 72s and order of presentation was counterbalanced.

### Stimuli

Milkshake stimuli were presented as two flavors, chocolate and strawberry. Chocolate milkshakes were made by combining 12 fl oz of whole milk with equal parts Garelick Farms Chug chocolate milkshake and Garelick Farms Chug cookies and cream milkshake. The strawberry milkshake was made by combining 32 fl oz of whole milk with 6 fl oz of Hershey’s strawberry syrup. One subject did not like the taste of strawberry and instead was given a vanilla milkshake created by mixing 32 fl oz whole milk with 6 fl oz of Eclipse vanilla syrup. Milkshakes were presented in an interleaved manner to reduce the possibility of sensory adaptation. Tasteless solutions were chosen individually by each participant, based on what they perceived “tastes most like nothing.” Tasteless solutions were one of either 25mM KCl /2.5mM NaH CO3, 18.75mM KCl/1.875mM NaHCO3, 12.5mM KCl/1.25mM NaHCO3, 6.25mM KCl/0.625mM NaHCO3, or 3.13mM KCl/0.313mM NaHCO3, dissolved in distilled water.

### Stimulus Delivery

0. 5 mL of milkshake was delivered into the mouth over 2s using an MRI-compatible gustometer system. The gustometer system *(51)* is a portable device controlling 11 programmable BS-8000 syringe pumps (Braintree scientific, Braintree, Ma). Pumps are loaded with 60 mL syringes filled with milkshake, which are infused into 25ft of Tygon beverage tubing (Saint-Gobain Performance Plastics, Akron, OH) connected a Teflon gustatory manifold on the MRI headcoil. The manifold consists of multiple channels that converge at a central point to allow delivery of a liquid stimulus to the tongue tip.

### Preprocessing

Neuroimaging data were preprocessed using SPM12 (Statistical Parametric Mapping, Wellcome Department of Imaging Neuroscience, London, UK) and BioImage Suite *(52)*. Participants’ functional images were corrected for geometric distortions resulting from susceptibility-induced field inhomogeneities. The FieldMap toolbox in SPM12 was used to calculate static distortions *(53)*. Corrections were then made for both static distortions and changes in these distortions resulting from head motion *(54, 55)*. Images were realigned, unwarped, and coregistered to each participant’s anatomical T1 image and then processed using a method combining segmentation, bias correction, and spatial normalization *(56)*. Functional images were slice-time corrected, normalized to a standard MNI template, and smoothed using a 6 mm FWHM isotropic Gaussian kernel. Normalized, motion-corrected images were further preprocessed using BioImage Suite. These additional steps included regression of the mean time courses of global signal, CSF, and white matter, in addition to removal of the linear trend and low-pass filtering.

### Construction of predictive networks

We performed Connectome-based Predictive Modeling (CPM; *14, 15, 18*) to determine whether task-based functional connectivity during milkshake receipt can predict individual differences in anthropometric measurements. To accomplish this, we first used a whole-brain functional atlas *(57)* to parcellate the brains of each subject into 268 nodes (i.e spatial ROIs) that maximize the similarity of voxel-wise time courses within each node. 268×268 connectivity matrices for each run were subsequently created in BioImage Suite by averaging the BOLD signal of each voxel within nodes *(52)*. Connectivity matrices were Fisher Transformed to convert the skewed distribution of Pearson r (or Spearman rho) values to an approximately normal distribution. Matrices were then averaged across runs to create a mean matrix for each subject, which were then collapsed across subjects to create 268×268×N-subject matrices.

In the N-1 training set, CPM uses a linear regression (i.e. Pearson’s r or Spearman’s Rho) to correlate each edge (i.e. connection) in each connectivity matrix to a behavioral measure of interest. Each edge, as a result, is then associated with a particular p-value and the most relevant edges are selected for further use via thresholding (P = 0.01). Edges that are positively correlated with behavior constitute a “positive prediction network,” while edges whose strengths are negatively correlated with behavior constitute a “negative prediction network.” Single-subject summary values are then calculated by separately summing the edge-strengths of positive and negative prediction networks *(15)*. These positive and negative “network strengths” are used in a linear model with behavior and applied prospectively to the left-out subject’s network strengths to generate predicted behavioral scores.

### Permutation Testing

To determine the significance of the obtained correlations between observed and predicted behavioral scores, we conducted 1000 repetitions of CPM using randomly shuffled observed scores to generate predicted scores and networks. The 1000 correlation coefficients obtained from permutation testing were used to comprise a null distribution against which the correlation coefficients in the Results section were tested for significance. Final p-values were calculated by determining the number of permutations, out of 1000, that generated correlation coefficients larger than those reported in Results.

### Visualization

Figures were constructed using BioImage Suite *(52)* and custom scripts in MATLAB^®^ 2016b. Anatomical locations were isolated using the Wake Forest PickAtlas Toolbox v. 3.0.5 (Wake Forest University, Winston-Salem, NC, USA) in SPM12 and the Automated Anatomical Labeling (AAL) atlas *(58)*.

## Code Availability

Matlab scripts to run the CPM analyses are available at (https://www.nitrc.org/projects/bioimagesuite/). Visualization tools are available online through BioImage Suite and accessible at (https://bisweb.yale.edu). Additional Matlab scripts for post-hoc analyses and visualization are available from the authors upon request.

## Data Availability

The data used to support the findings of this study are available from the corresponding authors (M.C.F. and D.M.S.) upon reasonable request.

## Acknowledgements

We would like to thank Corey L Horien and Abigail Greene for their help with data visualization. This work was supported by the National Institutes of Health training grant T32NS041228 (M.C.F.); the Yale Medical School Fellowship (M.C.F.); the National Institutes of Health R01 CA180030 (D.M.S.).

## Author Contributions

Conceptualization, D.M.S. and M.C.F.; Investigation D.M.S and M.C.F.; Software, R.T.C. and D.S.; Formal Analysis, M.C.F. and M.J.V.; Visualization, M.C.F. and M.J.V.; Writing – Original Draft, M.C.F. and M.J.V.; Writing – Review and Editing, M.C.F. and D.M.S.; Supervision, R.T.C., D.S., and D.M.S., Funding Acquisition, D.M.S.

## Competing Interests

The authors declare no competing interests.

