## Supplemental Table 1 for "Fingerprinting Adiposity and Metabolic Function in the Brains of Overweight and Obese Humans"

**Table S1. Participant Characteristics of Full Sample.**

| Variable of Interest | Mean ± Standard Deviation |
|---|---|
| Age | 30.7 ± 6.57 |
| Sex | 50 male, 17 female |
| BMI | 33.3 ± 6.52 |
| BF% | 40.2 ± 8.61 |
| Waist Circumference | 38.2 ± 6.21 |
| Waist/Hip Ratio | 0.9 ± 0.09 |
| Fasting Glucose (n=67) | 96.6 ± 13.98 |
| Fasting Insulin (n=59) | 18.4 ± 16.01 |
| HOMA-IR (n=59) | 4.3 ± 3.96 |
